# Generation of Bessel beam lattices by a single metasurface for neuronal activity recording in zebrafish larva

**DOI:** 10.1101/2023.02.12.528189

**Authors:** Anna Archetti, Matteo Bruzzone, Giulia Tagliabue, Marco dal Maschio

## Abstract

Bessel Beams (BBs) and BB lattices are structured-light excitation profiles frequently applied in material processing, nonlinear spectroscopy and in many fluorescence microscopy methods such as Light Sheet Microscopy (LSM). In LSM, BBs and BB-lattices offer wider excitation profiles, higher acquisition rate, enhanced resolution, and improved signal-to-noise ratio, while reducing the overall phototoxicity. However, this performance improvement typically comes at the cost of layout complexity and spatial constraints, originating from the optical arrangement required for obtaining BB features and for multiplexing the BB in a lattice of beamlets. Here, we introduce a novel method for encoding in a single flat element all the optical operations required to generate a BB lattice, including those of the excitation objective. We assessed the effective capabilities of this approach, using Meta-Surface (MS) technology to fabricate the corresponding flat optical element and to characterize its optical figures. Finally, we demonstrated its actual application in LSM, recording neuronal activity at cellular resolution in the zebrafish larval brain using fluorescence based neuronal activity reporters. In perspective, this approach, applied here for LSM, prompts a step forward in the BB versatility and in the BB application scenarios.

## Introduction

High-resolution fluorescence imaging is a fundamental approach for structural and functional non-invasive studies in life science. Under the increasing need for high acquisition rates and large volumes of investigation, many techniques have been developed. Point scanning methods, such as confocal and multiphoton-photon microscopy, present high signal-to-noise ratios and the investigation depth within the tissue. Yet they are intrinsically limited by the scanning process that has to balance the scanning rate with the amount of signal integration. On the other hand, parallel approaches, like light sheet microscopy (LSM), when the tissue scattering is not a constraining factor, combine simultaneous excitation with one-shot acquisition of the entire Field of View (FoV) in a few milliseconds. This method ensures at the same time high signal collection efficiency, reduced out-of-plane excitation, and low photodamage^1^. In an LSM system, the sample is typically excited with a few micrometers thick sheet of light and the signal is collected at once from the entire FoV along the direction orthogonal to the illuminated area. A suitable excitation beam, with a flattened profile in the direction perpendicular to the light propagation direction, is typically achieved by collimating a Gaussian beam onto a cylindrical lens^1^, or by using the diffraction of a beam entering the sample glass slide at high angles through an objective^2–5^ or a prism^6^. In other common solutions, usually referred to as digitally scanned light-sheet microscopy (DSLM)^7^, a virtual light sheet is obtained by scanning in the transversal direction a beam across the sample. Along with the traditional use of a single Gaussian beam, more recently, approaches based on multiple Bessel Beams (BBs), such as lattice light sheet (LLS)^8–10^ and universal lattice light sheet^11^, have been developed. With respect to Gaussian beam, BBs present a strongly elongated point spread function along the propagation direction and a typical cross section with a narrower central peak^12,13^. Such narrow intensity profile, invariant over larger propagation lengths and more resilient to diffraction, renders Bessel beams ideal for optical approaches with improved resolution, excitation profile extension, and reduced phototoxicity.

However, generation of multiple BBs typically require specialized optical elements that tend to render the optical layout rather complex, spatially extended and articulated. In fact, moving from Gaussian to BB-based excitation calls in for the integration of wavefront modulation components, e.g. an axicon phase plate or an apodization filter (annular mask)^13^, to render a more elongated excitation profile. Furthermore, generating multiple BBs at the sample, like in LLS, frequently requires beam-multiplexing elements, like Spatial Light Modulators (SLMs)^8^ or other Diffractive Optical Elements (DOEs)^9^ for generating the light sheet. Ultimately, these two optical components need to be integrated at different positions along the optical path so to have a proper optical conjugation with the excitation objective. Here, we introduce a novel solution for generating excitation profiles with multiple BBs by means of a single flat optical element, overcoming the complexity of the current designs. We developed a method to encode on a single optical surface all the functions of the excitation objective, of the apodization mask for the generation of a BB profile, and of the beam multiplexing, so to render, upon light propagation, a lattice with multiple beamlets at the sample. To test this approach, we designed and fabricated an optical component using Meta-Surface (MS) technology. MSs are engineered surfaces patterned with nanostructures to control different light properties, like amplitude, phase, and polarization with sub-wavelength resolution^14^. Following a characterization step confirming the generation of the expected BB patterns, we moved further and aligned the engineered MS along the excitation path of a LSM microscope. Using this MS-based LSM configuration, we recorded neuronal activity from the brain of a zebrafish larva with a spatial resolution sufficient to resolve the different cells.

## Results

### Generation of a single BB by wavefront modulation on a single surface

The electromagnetic field of a Bessel beam presents an intensity profile extended along the *Z*-propagation direction with an almost uniform *XY*-cross-section showing a central maximum surrounded by a series of side lobes of decreasing intensity. It is known that the spectrum of this field recapitulates a ring-delta function, with wavevectors lying on the surface of a cone^12,13^. Thus, confining the field intensity of a Gaussian beam to a thin ring in the source space, it renders a Bessel beam in the Fourier space of a converging lens or objective. This principle, already introduced as alternative to axicon-based approaches, adopts an apodization filter, to select a ring-shaped portion of the original beam, and its conjugation by a 4f optical arrangement to the back focal plane of an objective to render a Bessel beam at the sample. We reasoned that, in principle, one can collapse this complete optical path and encode on the same optical element the functions of the ring and of the objective, so to build upon light propagation the convolution of an annular intensity profile and a wavefront phase modulation corresponding to a focusing lens (**Figure 1a** and **b**).

**Figure 1.**
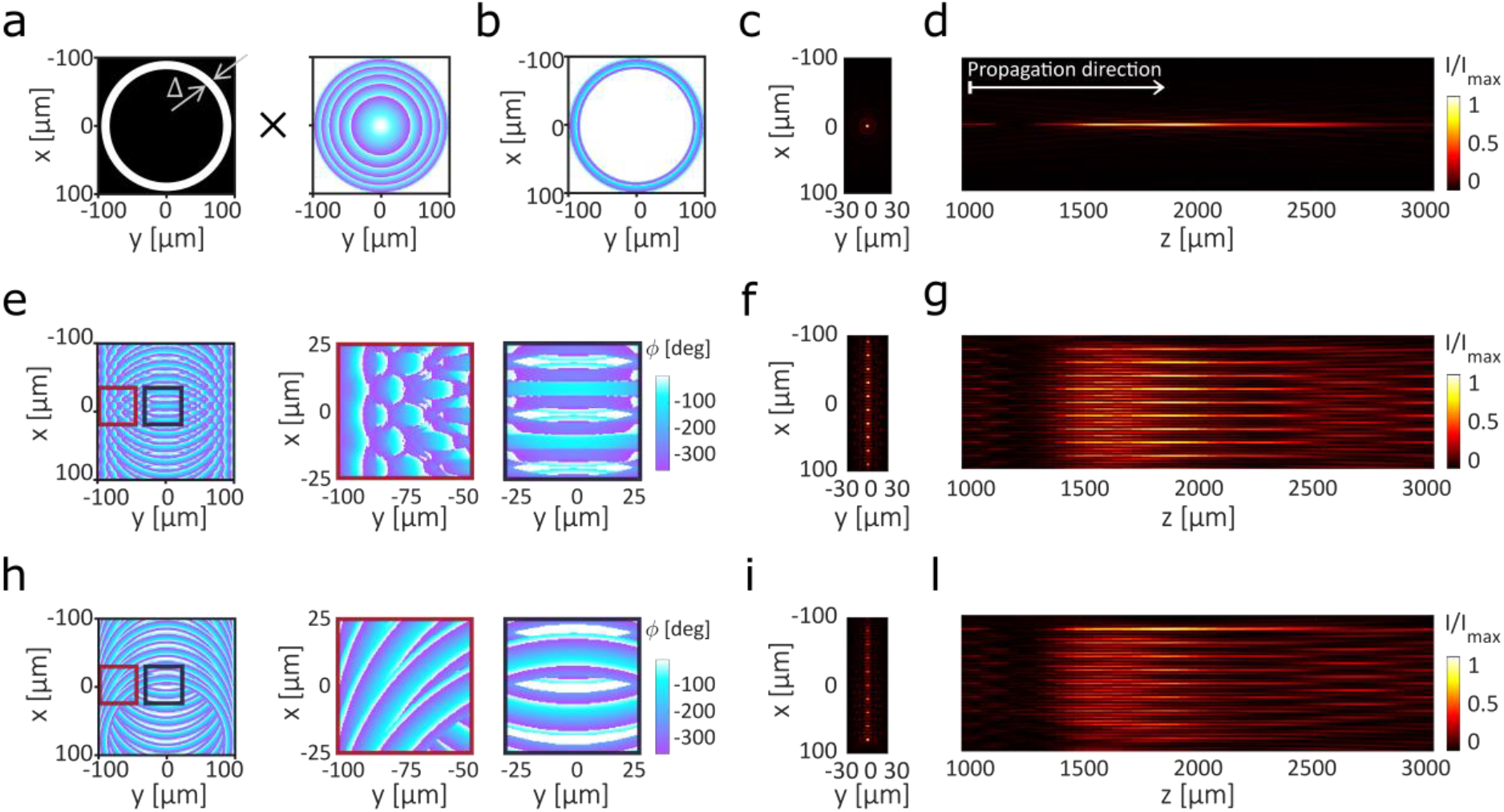
Working principle of the proposed single element Bessel Beam Lattice excitation. **a**, The amplitude annular mask (**a-left**) with outer radius *R* = 100 μ*m*, and inner radius *R* − ∆ = 83 μ*m* is combined with the phase profile of a convergent lens with focal length *f* = 2000 μ*m* (**a-right**) to produce a phase profile with a ring shape **b**, Ring-phase mask obtained from the operations described in (**a**) for the generation of a single BB **c**, BB intensity profile in the XY plane transversal to the beam propagation direction; **d**, BB intensity profile in the beam propagation plane XZ. **e**, Total BB phase profile *ϕ*_***TOT***_ retrieved *with method-1* (**e left**). Highlighted regions in **e-left** show the phase profile where the rings overlap on the left (**e middle**) and in the middle (**e right**). **f and g**, Simulations of the BB intensity profiles generated with the phase mask *ϕ*_*j*_ described in **e. f**, intensity profile of the BB in the XY plane; **g**, intensity profile of the BB in the XZ plane. **h**, The BB phase profile *ϕ*_***TOT***_ generated *with method-2*. Highlighted regions in **h left** show the phase profile where the rings overlap at the left (**h middle**) and in the middle (**h right**). **i and l**, Simulations of the BB intensity profiles generated from the phase mask shown in **h. i**, intensity profile of the BB in the XY plane; **l**, intensity profile of the BB in the XZ plane.

Assuming a planar wave at the input, a phase mask 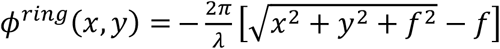, defined on a spatial domain in the XY source plane with 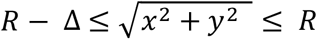and zero elsewhere, works at the same time as an optical Fourier transformer characterized by a focal length *f* and an amplitude annular mask parametrized by an external radius *R* and a thickness ∆ (**Figure 1a**). According to our simulations, the expected intensity profile at the sample (**Figure 1c** and **d**) resembles a BB whose spatial properties depend on the design parameters *R*, ∆ and *f* (**Figure 1a**). *R* and *f* allow for controlling the effective NA of the beam and its longitudinal position; the annulus ∆ impacts on the Gaussian-over-Bessel feature balance. Indeed, a beam generated by an annulus mask with finite aperture, will have a mix of characteristics of both Gaussian and Bessel beams. The Gaussian envelope contributes in suppressing the side lobes of the BB (due to the continuum of wavevectors across the annulus)^8^. On the other hand, the elimination of wavevectors inside the inner diameter of the annulus, ensures a much longer depth of focus than a traditional Gaussian beam. Therefore, the thinner the annulus ∆, the more prominent the Bessel contribution: the central lobe of the beam becomes thinner and the longitudinal dimension of the illumination profile more extended.

### Generation of multiple BBs by wave-front modulation on a single surface

It is common to use BB in a lattice-based illumination scheme^8–10^, with multiple beams illuminating the sample. Optical lattices are periodic interference patterns, organized in 1D or 2D regular arrays of beamlets, generated by the coherent superposition of a finite number of plane waves^15^. Interestingly, we noticed that it is possible to extend our approach and to encode on the same plane a total phase mask *ϕ*_***TOT***_ capable of generating multiple BBs and thus, as previously reported, a BB lattice accordingly to any desired 3D geometry. We identified two approaches for obtaining the required phase profile. In the first case, (*method-1*), the total phase profile *ϕ*_***TOT***_ is the argument of the sum of the fields *E*_*j*_ each corresponding to the *j-th* Bessel beam (**Figure 1e-g**): 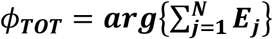 where 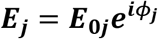. Each field *E*_*j*_ is characterized by a phase*ϕ*_*j*_(*x, y*) = *ϕ*^*ring*^(*x* − *jP, y*), replica at a x-distance *jP* of the original phase mask described above, and an amplitude

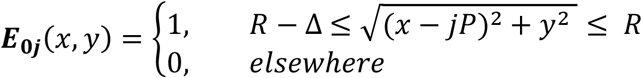

representing the P-shifted annular mask. In the second approach (*method-2*), the total phase profile *ϕ*_***TOT***_ of 1D lattice of BBs is generated by overwriting each consecutive field 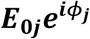 (**Figure 1h-l** and **Supplementary Note 1**).

It is known that the period *P* separating the beams can be optimized 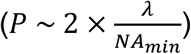 such that to produce a destructive interference between the side lobes of adjacent beams and to maximize the contribution of the main lobe of the Bessel beams^8^. With the design parameters adopted for the ring properties (outside ring radius *R* = 100 *μm*, inner radius *R* − ∆= 83 *μm*, focal length *f* = 2000 *μm*, and numerical aperture relative to the inner diameter of the annulus *NA*_*min*_ = 0.038), we could obtain Bessel beams laterally shifted by *P* ~ 20 *μ*m, (**Figure 1h-l**). To test the outcome of the developed methods, we implemented a beam propagation algorithm for calculating the spatial intensity profiles of the generated BBs (**Methods**Error! Reference source not found., **Supplementary Note 2**). This analysis confirmed that such single plane wavefront modulation patterns render upon light propagation multiple BBs according to the design parameters: each beam with a Full With at Half Maximum (FWHM) of *FWHMx* ~ 4 *μm* and *FWHMy* ~ 7 *μm* on the transversal X and Y-direction and *FWHMz* ~ 700 *μm* along the beam propagation direction.

### Generation of BB lattices using Meta-Surface technology

To assess the effective capabilities of a flat optical element with the features described above, we proceeded with its fabrication adopting Meta-Surface (MS) technology. This approach uses arrays of nanopillars few hundreds of nanometer high to control the properties of the electromagnetic field^16–19^. MSs can be used to simultaneously control the amplitude, the phase and the polarization of a propagating light wavefront, thus enabling color routing^20,21^, polarization-multiplexing^22–24^, and focusing^25^. We considered a set of possible materials for the fabrication and selected Silicon Nitride (SiN^x^). SiN^x^ is a dielectric material compatible with CMOS fabrication processes^26,27^, with good chemical and thermal stability^28,29^, with high index contrast compared to the oxide cladding (∆*n*~0.5) and with a broadband transmission extended to the visible spectrum^26,30^. Therefore, we fabricated and tested a series of polarization-insensitive SiN_x_ Huygens MSs designed to operate at *λ* = 478 *nm* for producing multiple BBs (**Figure 2a**).

**Figure 2.**
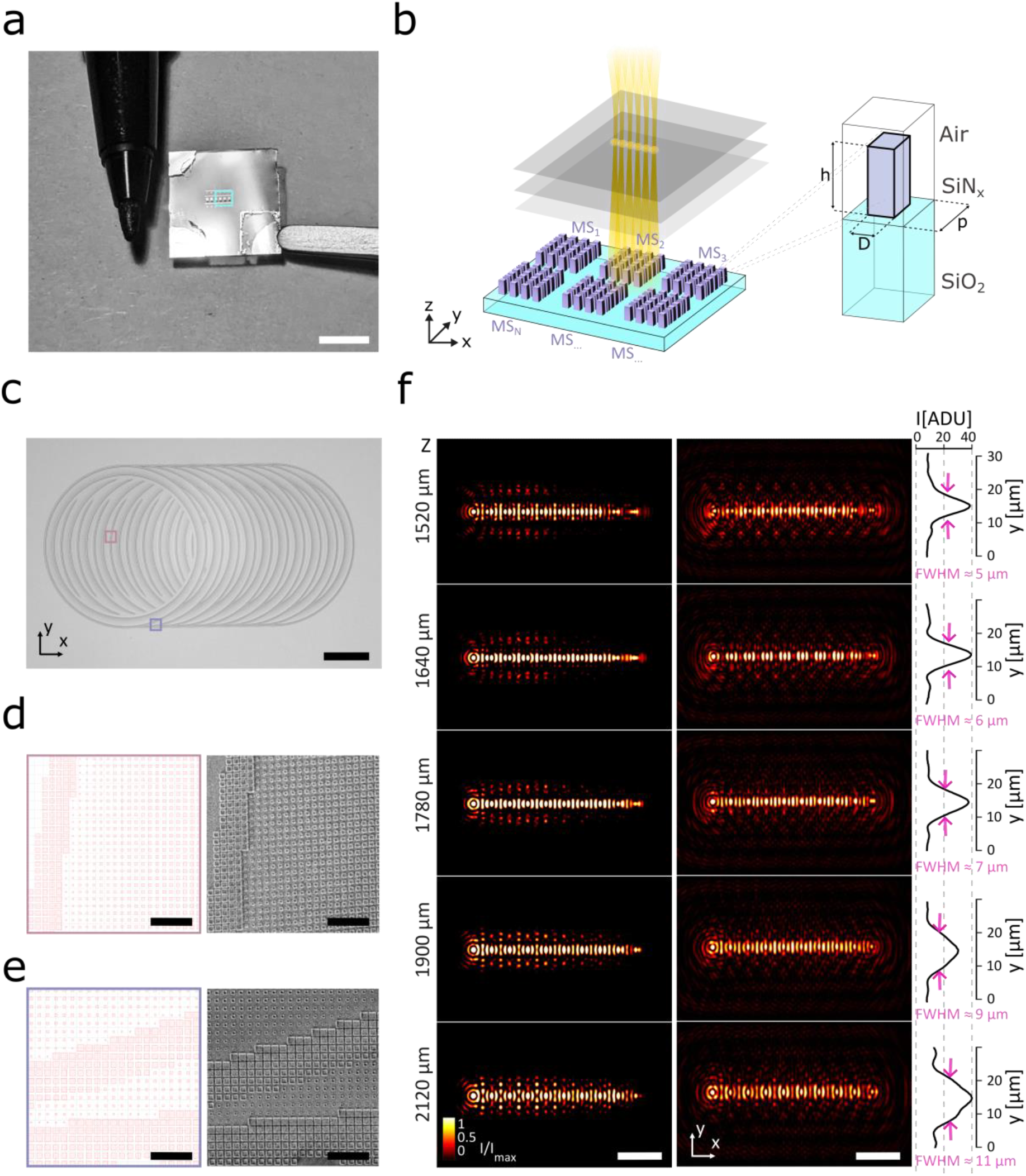
Single metasurface based generation of BB lattice. **a**, Exemplar of one fabricated chip with the metasurfaces (MSs). The MSs structures appear in reflectance as light-grey stripes on the chip surface. **b** Schematic of the unit nano-pillar of the metasurfaces (MSs). **c**, An optical wide field image of one of the metasurfaces for Bessel beam lattice light sheet (BB-LLS) excitation. The MS shows the characteristic rings of our BB-LLS principle: each ring simultaneously behaves as both converging lens and annular mask and thus transforming each quasi-Delta-Ring into a quasi-Bessel beam. **d** Magnified view of marked central region in (**c**): MS layout (**left**) and corresponding scanning electron microscopy (SEM) image (**right**). **e**, Magnified view of marked peripheral region in (**c**): MS layout (**left**) and corresponding SEM image (**right**). The SEM images show the nanometric scaling of the square nano-scatter cross-section. **f**, Simulated beam profiles of the BB-LLS generated by our ML layout at different propagation depth (**left**); experimentally measured beam profiles of the BB-LLS generated by our fabricated ML (right) and their corresponding intensity profiles along the Y axis. Experimentally measured full width at high maximum (FWHM) of the BB-LLS Y-profiles of < 7 µm. Scale bars: 0.5 cm (**a**), 50 µm (**c** and **f**), 2 µm (**d** and **e**).

Our MS design presents a series of nanopillars, built on top of Silicon dioxide substrate (**Figure 2b**), with a different lateral dimension to locally modulate the phase of the incoming wavefront, hence acting as a phase-delay element. We used finite difference time-domain (FDTD) numerical simulations (COMSOL Multiphysics®, **Methods** and **Supplementary Note 2**) to study the electromagnetic field modulation of the nanopillar and identified a library of 28 structures ensuring the coverage of the phase range *ϕ* = [0, 2π] with a transmittance efficiency over 50% (**Supplementary Figure 1**). These nanopillars present a square cross section with a variable side length D, ranging between 60 and 340 nm at steps of 10 nm, and with a fixed height *h* = 520 nm (**Figure 2b**). The designed MS active area resulted about 200 µm in diameter with nanopillars distributed with a fixed spatial period (*p* = 400 nm). Then, we developed a MATLAB routine to convert the continuous phase profile obtained with the method-2 to the discrete phase map of the MS (**Supplementary Note 3, Supplementary Figure 2**, and **Supplementary Figure 3**). The numerical simulations confirmed that the adopted MS layout generates a set of ten Bessel beams in close agreement with the theoretical model (**Supplementary Figure 4**).

Based on this confirmation, we fabricated a set of MSs (MS_1_, MS_2_, …, MS_N_) by ebeam lithography according to a process flow we developed (see **Figure 2c-e, Methods, Supplementary Note 4**, and **Supplementary Figure 5-8**). The characterization of the intensity profile generated by the MS confirmed that we obtained a set of ten BBs with ∼ 4 × 7 × 700 µm_3_ (FWHM X, Y, Z) size and extending over a lateral dimension X of about 200 µm, close to the values expected from the simulations (**Figure 2f, Supplementary Video 1**, and **Supplementary Figure 9**).

### Metasurface-based lattice light sheet imaging of zebrafish neuronal activity

We integrated the fabricated MS into the excitation path of a custom LSM setup to demonstrate the performance of our approach. The MS was inserted between the laser source and the sample, replacing the optics and the objective of the typical LSM excitation optical path. We tested our system by recording neuronal activity from zebrafish larvae expressing the genetically encoded activity indicator GCaMP6s in the majority of the neurons in the brain. This light-based reporter, changing its fluorescence emission as a function of the neuron action potential firing rate, allows to reconstruct the spatial pattern of the neuronal activity at cellular resolution. For recording brain activity, the MS was aligned parallel to the sagittal plane of the zebrafish brain (see **Figure 3a** and **Methods**), so to generate BB lattice excitation on a large area of the brain along the medio-lateral direction. By means of an orthogonal detection arm equipped with a CMOS camera, we successfully recorded neuronal activity in multiple brain regions (**Figure 3b**) characterized by different amount of light scattering, like the Pallium (**Figure 3d**), Tectum Opticum and Cerebellum (**Figure 3e**) and Medulla Oblungata (**Figure 3f**). Zebrafish brain presents cells with 6-7 µm typical diameter and a degree of neuronal packing with almost no extracellular space. Even in these challenging conditions, the quality of the imaging allowed us to segment the area of the individual neurons by means of an automated segmentation analysis pipeline commonly used for this application^31^ (**Figure 4g-i** and **Methods**). Temporal series of the neuronal firing associated with spontaneous activity in the brain were easily retrieved, presenting broad spatial patterns of activity involving large population of cell interleaved with segregated clusters of activity in small neuronal subsets (**Figure 4l-m**). Thus, the obtained dataset confirmed the possibility to use our BB lattice generation method on a challenging application, like fluorescence-based functional brain imaging using LSM, typically characterized by articulated arrangements.

**Figure 3.**
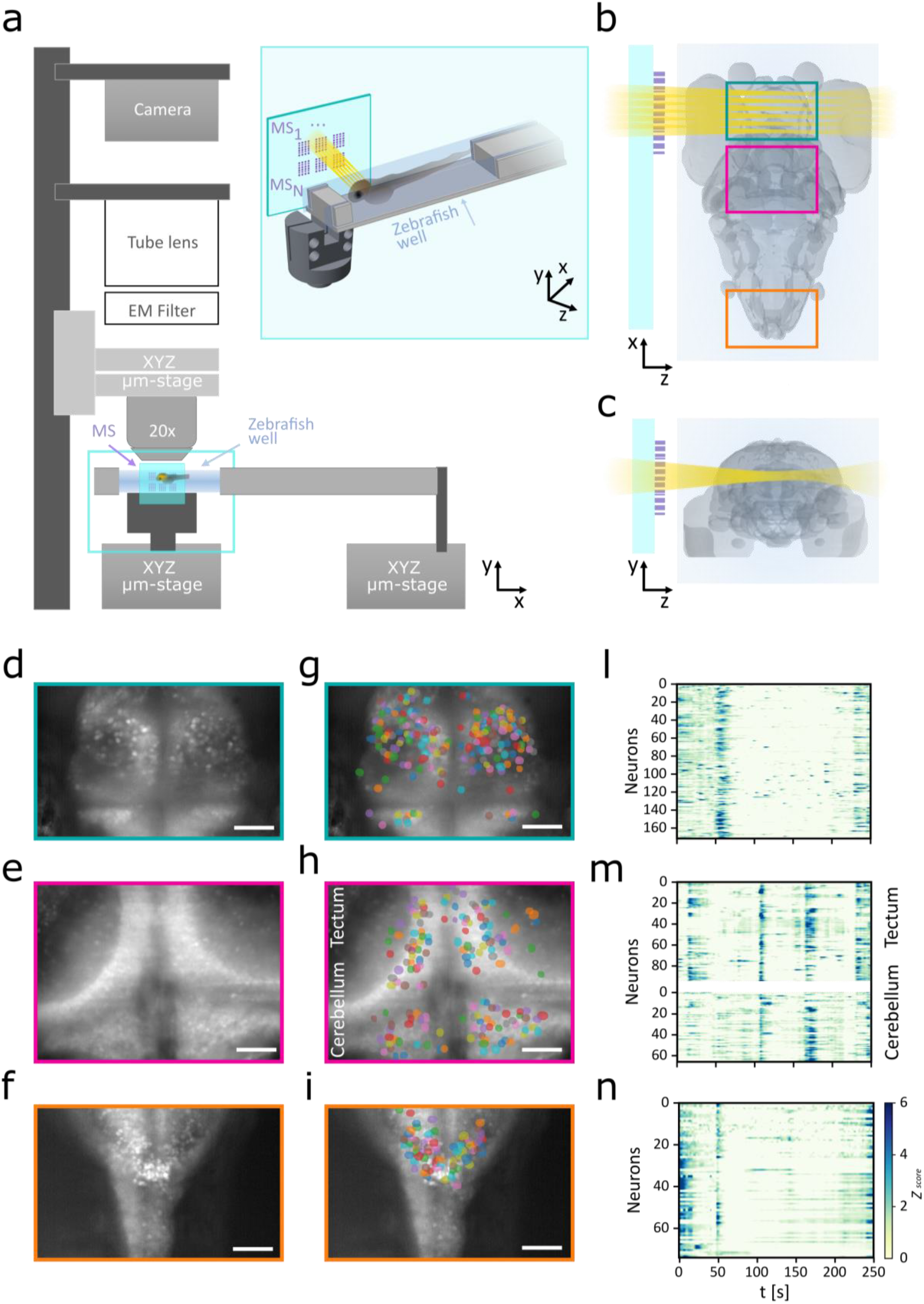
MS-based Bessel beam lattice light sheet imaging of the zebrafish brain activity. **a**, MS-based BB LLS microscope. The setup is made by two independent arms: the horizontal illumination modulus, where the BB light sheet is generated by the metasurface (MS) system and the vertical imaging column, where the fluorescence emission signal is collected by a custom made up-right microscope. The zebrafish is embedded in agarose inside a millimetric 3D printed well. The MS chip is mounted parallel to the sagittal plane of the zebrafish with a custom-made precision holder (cyan insert in **a**). **b**, Horizontal projection of the zebrafish larva brain. The colored boxes refer to the areas imaged with the MS-based LLS microscope. **c**, Coronal projection of the zebrafish larva brain. **d-f**, Maximum intensity projection of the brain regions highlighted in **b. g-i**, Segmented neurons extracted from the zebrafish brain regions highlighted in **b. l**, Raster plots showing the calcium activity profiles of the identified neurons of the region shown in **g. m**, Raster plots showing the calcium activity profiles of the identified neurons of the region shown in **h. n**, Raster plots showing the calcium activity profiles of the identified neurons in region shown in **i**. Scale bar: 50 µm (**d-i)**

## Discussion and Conclusions

Adopting methods of wavefront engineering, we present here a method to integrate in a single plane all the optical manipulations necessary to generate at the sample an excitation profile with a lattice of Bessel beams. Applying this strategy and using MS technology, we fabricated the corresponding optical element with a total area smaller than one by one-millimeter square and with an active area of 200 µm in diameter. Its integration along an optical path allowed us to validate the fabricated component in a challenging application, the recording of the neuronal activity from the zebrafish larva brain at cellular resolution. With respect to the current state of the art, the proposed approach presents a number of advantages. First, collapsing all the optical operations required for the generation of BB lattice in one single optical element, we show that is possible to substantially reduce the footprint and the complexity of the traditional microscopy configurations used for this purpose. This has the potential to extend further the level of integration and miniaturization that an optical system could achieve. Second, taking advantage of the “in-plane” beam multiplexing, our approach allows for maximizing the light utilization efficiency. Indeed, traditional LLS designs block the input beam light at the center of the apodization mask, while, in our design, the central part of the input beam contributes to the generation of other BBs in the lattice. Moreover, generating the BB lattice with a set of independent phase modulation rings, the focal position and the depth of focus of each BB and their numbers can be independently designed, potentially enabling the generation of envelopes of BBs with an arbitrary 3D structure along the light propagation direction and arbitrary extension in the perpendicular plane, and so overcoming the limitation of the regular lattices. The single plane strategy we devised for the generation of BB lattice could be in principle embodied using different technologies for the fabrication of flat optics, like traditional Fresnel Lenses or DOEs. We adopted here MS technology as it supports the simultaneous but independent modulation of different parameters of the electromagnetic field, like amplitude, polarization and phase. Moreover, another key advantage of MS technology derives from their potential tunability and thus from their intrinsic capability to dynamically adapt to different optical requirements. Noteworthy, the sub-millimetric size of the optical elements designed according to the proposed strategy allow for the fabrication on the same substrate of multiple lens layouts, each with optical properties tailored around different requirements, making the fabricated optical element a flexible and versatile multi-scale and multipurpose imaging system. In conclusion, our design, opens a range of possible application scenarios of BB lattice not only in microscopy development but more in general in all those fields where small-footprint methods for light shaping with multiple BBs is instrumental for increasing the parallelization, the throughput and resolution levels.

## Methods

### Numerical simulations of the nano-pillars electromagnetic response

The electromagnetic response of individual nano-scatter must be engineered such that their phase parameter space satisfies the target phase *ϕ*(*x, y*) within the selected tolerance error - fixed here at ±2 degrees in our design. To predict the phase accumulated by each nano-pillar and their transmittance we used COMSOL Multiphysics® platform which is widely used for finite difference time-domain (FDTD) numerical simulations of optoelectronic and photonics devices. The results of transmittance and phase of each nano-pillar are described in **Supplementary Note 2** and shown in **Supplementary Figure 1**.

### Metasurface design and fabrication

To design our metasurface and create the layout required for the fabrication, we implemented a semi-automatized MATLAB routine (see **Supplementary Note 3, Supplementary Figure 2** and **Supplementary Figure 3** for further details).

We fabricated the SiN_x_ metasurfaces at the EPFL Center of Micro-Nanotechnology (CMi). In the last years, these features have made SiN_x_ manufacturing technology more and more accessible for micro-nanofabrication of optical components for imaging application^32,33^.

First, a 520nm-thick silicon nitride (SiN_x_) layer is deposited on the silicon dioxide (SiO_2_ - 525 µm thick) as core material. Secondly, to create the MS pattern, we spatter 30nm of chromium (Cr) as hard mask and as opaque reflective layer for a proper focusing of the ebeam-tool and we spin coat 200 nm of hydrogen silsesquioxane (HSQ) as positive resist for the e-beam lithography. The MS pattern is thus created on the photoresist after development. The MS structure is then transferred into the Cr and SiN_x_ layer by one first step of ion beam etching (IBE) followed by a high-density inductively coupled plasma (ICP) etching step based on fluorine chemistry (CHF_3_/SF_6_). Finally, the residual of Cr and photoresist is stripped by wet acid etching. The main fabrication steps are depicted in **Supplementary Note 4** and **Supplementary Figure 5**. Images of the fabricated chips are shown in **Figure 2c-e, Supplementary Figure 6**, and **Supplementary Figure 7**.

### Metasurface-based Bessel beam characterization

The profile of the Bessel beam lattice light sheet was measured by acquiring its intensity along the propagation direction. For these characterization measurements, a 20x Olympus objective was mounted with a kinematic mount on a one-axis motorized stage (PT1/M-Z8, Thorlabs). The objective was oriented parallel to the propagation beam and adjusted such to focus on the chip surface that we defined as the zero-propagation-position, z = 0 µm (**Supplementary Figure 8**). The metasurface chip was fixed in the vertical slit of a custom-made rotating holder mounted on a three-axis stage to align is surface perpendicular to the incoming beam. The BB light sheet profiles were acquired at different propagation positions z, from z_0_ = 500 µm up to z = 3500 µm with a step size of z_step_ = 5 µm with a CMOS camera (XIMEA, MQ013RG-ON) at an exposure time of 1 ms. All the metasurface characterization measurements was performed with a 488 nm Laser (Cobalt 06-01) at a laser power value of Pw ∼ 0.3 mW. The BB-LLS thickness was then estimated by measuring the full width at half maximum (FWHM) of the X and Y intensity profile at a depth ranging from 1500 µm up to 2200 µm every 120 µm.

### Metasurface-based BB-LLS microscope

The metasurface chip for BB-LLS illumination is integrated into a custom up-right microscope thanks to two mechanical holder we designed to fit the required physical constrains imposed by the MS chip, the sample holder, and the imaging objective (**Figure 3a, Supplementary Figure 10 and Supplementary Figure 11**). The MS chip holder can be easily placed into the XYZ-stage (a three-axis stage, MBT616D/M, Thorlabs) to enable the aliment of the MS with the input beam (488 nm Cobolt 06-01 Laser). The sample holder is mounted on a second identical stage to adjust the sample sagittal plane parallel to the chip surface. Emitted light from the sample was collected by the air objective lens (MPLAN APO20 20x/NA0.42 Mitutoyo), then imaged by a tube lens (fTL = 200 mm) onto the CMOS camera (XIMEA, MQ013RG-ON).

### Sample Preparation for live imaging

Zebrafish brain imaging experiments were carried out in line with the current Italian legislation (Decreto Legislativo 4 Marzo 2014, n.26) and were approved by the Ethical Committee of the University of Padua (61/2020_dalMaschio) and adhere to the ARRIVE (Animal Re-search: Reporting of In Vivo Experiments) guidelines.

Larvae were raised at 28°C on a 12h light/12h dark cycle using standard procedures. Elavl3:H2B-GCaMP6s larvae were used for brain imaging. Five days post-fertilization (dpf) larvae were embedded in 2% low melting point agarose gel on a 3D printed custom-made support (**Supplementary Figure 10**).

### Zebrafish brain imaging and data analysis

The microscope and all components were controlled with µManager^34^. The 488 nm laser output power was set to deliver about 1 mW at the sample. The brain of the embedded living zebrafish was moved into focus using the coarse and fine focusing screws of the translation stage. The spatial and temporal resolution of the image stack acquisition were 0.240 µm/pixel and 500 ms/frame, respectively. Spontaneous activity was recorded for 250 seconds from different brain regions and with different subjects. The recorded raw movies of the brain regions were subsequentially processed with Suite2p^31^ for automatic segmentation of the active neurons and to extract their calcium activities.

## DATA AVAILABILITY

All data and software used to support the results of this manuscript are available from the Lead Contact upon reasonable request.

Original data is available on https://doi.org/10.5281/zenodo.7589979

## ACKNOWLEDGEMENTS

We thank Devis Pantano from the Physics Department of Padua for his support in the design and fabrication of the sample holders; we thank Marco Salamanca for support in the zebrafish preparation. Metasurface chips were fabricated at the EPFL Center of Micronanotechnology (CMi). This work was funded by European Union’s Horizon 2020 research and innovation program under the Marie Sklodowska-Curie grant agreement No. 898315 (FLAMMES).

