## Supplementary files for "Generation of Bessel beam lattices by a single metasurface for neuronal activity recording in zebrafish larva"

[3]*École Polytechnique Fédérale de Lausanne (EPFL), 1015 Lausanne, Switzerland*

**This file includes:**

- Supplementary Notes 1-4
- Supplementary Figures 1-11
- Supplementary Table 1-2,
- Supplementary References

**Other supplementary material includes:**

Videos from S1 to S3

### SUPPLEMENTARY NOTES

#### *S1. Design of the phase profile of a Bessel beam array*

The first approach (method-1) for the generation of the phase profile of a Bessel beam array relies on the following main steps.

**Step (1)**: the phase profile $\phi_j$ of a convergent lens with the desired radius $R_j$ and focal length $f_j$ is defined by

$$\phi_j(x,y) = -\frac{2\pi}{\lambda}\left[\sqrt{\left(x-P_j\right)^2+\left(y-Q_j\right)^2+{f_j}^2}-f_j\right];\ x,y \in \mathbb{R}^2 : x,y \leq\ R_j$$

where j is an integer number: $\mathrm{j} = 1,2,..\mathrm{N}$ to index the beam and $N$ is the desired total number of beams in the array. $P_j$ and $Q_j$ are two spatial parameters defining the position of the beam -j. The values of $R_j$, and $f_j$ set the desired focal position and the effective numerical aperture ($NA_j$) and therefore the maximum beam thickness and the minimum depth of focus of the beam *j*.

**Step (2)**: the mask $M_j(x,y)$ is created as a round iris. The mask is defined by:

$$M_j(x,y) = \begin{cases} 1, & R_j - \Delta_j \leq \sqrt{(x-P_j)^2+(y-Q_j)^2}\ \leq\ R_j \\ 0, & elsewhere \end{cases}$$

with external diameter $R_j$ and thickness $\Delta_j$. The radius of the ring must have the same value of the radius defined for $\phi_j(x,y)$ in step 1.

The thinner is the thickness, smaller $\Delta_j$, lower the beam thickness and longer its depth of focus.

**Step (3)**: the phase profile $\phi_j$ is multiplies by the mask $M_j$ such to obtain the phase profile:

$$\phi_{ring,j}\,(x,y) = \begin{cases} -\frac{2\pi}{\lambda}\left[\sqrt{(x-P_j)^2+(y-Q_j)^2+{f_j}^2}-f_j\right], & R_j-\Delta_j \le \sqrt{(x-P_j)^2+(y-Q_j)^2} \le R_j \\ 0, & elsewhere \end{cases}$$

**Step (4)** the field distribution is $\boldsymbol{E_j} = \boldsymbol{E_{0j}}\boldsymbol{e}^{\boldsymbol{i}\phi_{ring,j}}$ of an electromagnetic wave characterized by a phase profile $\phi_{ring,j}$ equal to the phase profile obtained in step (3) and with amplitude $E_{\mathbf{0},\boldsymbol{j}}$**:**

$$E_{0j}(x,y) = \begin{cases} 1, & R_j-\Delta_j \le \sqrt{(x-P_j)^2+(y-Q_j)^2} \le R_j \\ 0, & elsewhere \end{cases}$$

**Step (5)**: **Step (1)** to **Step (4)** are repeated for *N* times corresponding to the desired number *N* of beams in the array. At each iteration *j*, the phase profile can be designed with the desired focal length $f_j$, ring thickness $\Delta_j$ and ring radius $R_j$. The ring position can be shifted of the desired parameters $P_j$ and $Q_j$.

**Step (6)**: the total phase profile $\phi_{TOT}$ is computed as the result of the argument of the sum of the fields $E_{\boldsymbol{j}}$ each corresponding to one Bessel beam shifted in space of the desired parameters $(P_j, Q_j)$. Therefore, the final phase profile is $\phi_{TOT} = arg\{\sum_{j=1}^{N} E_j\}$ where $\mathrm{j} = 1, 2, ..N$ and $N$ is the total number of rings.

The second approach (method-2) for the generation of the phase profile of a Bessel beam array relies on the following main steps:

**Step (1)** to **Step (3)** the same as in method-1.

**Step (4)**: step (1) to step (3) are repeated for *N* times corresponding to the desired number *N* of beams in the array. At each iteration *j*, the phase profile $\phi_{ring,j}$ can be designed with the desired focal length $f_j$, ring thickness $\Delta_j$ and ring radius $R_j$. The ring position can be shifted of the desired parameters $P_j$ and $Q_j$.

**Step (5):** the values of the total phase profile $\phi_{TOT}(x,y)$ are the values of each single ring $\phi_{ring,j}$ and in all points $(x,y)$ belonging to multiple rings, $\phi_{TOT}(x,y)$ always takes the value of the phase of the ring $\phi_{ring,j}$ with minimum index.

Therefore, in this case the total phase profile $\phi_{TOT}(x,y)$ is defined as:

$$\phi_{TOT}(x,y) = \phi_{ring,j}(x,y)$$

Where j is the natural number minor or equal to N, defined by

$$j(x,y) = \min k \,:\, R_k - \Delta_k \leq \sqrt{(x-P_k)^2 + (y-Q_k)^2} \leq R_k$$

***S2. Numerical simulation for the study of the nano-pillars response***

To engineer the nano-pillars and to extract their transmittance and phase, we studied their electromagnetic response with COMSOL Multiphysics module (version 5.5) which solves the Maxwell's equation:

$$\nabla \times (\nabla \times \boldsymbol{E}) - k_0^2 \epsilon_r \boldsymbol{E} = \boldsymbol{0} \qquad (1)$$

where $\epsilon_r = (\mathrm{n} - i\mathrm{k})^2 = n^2$ is the relative permittivity, $\mathrm{n}$ and $\mathrm{k}$ are the real and the imaginary part of the refractive index $n$ respectively, $k_0 = 2\pi/\lambda$ is the wave number of free space. Each silicon nitride nano-pillar unit was simulated with a periodic boundary condition along the transverse direction with respect to the propagation of light and a perfectly matched layer and input/output ports boundary conditions along the longitudinal direction. We then extracted their transmission $T$ and phase $\phi$ of the electric field from the scattering parameter $S_{21}$ of the $S - parameter$ matrix, measured from the eigenmode expansion of the electromagnetic field at the output port 2 (see **Supplementary Figure 2a**). Therefore, if $E_1$, $E_2$, $E_3$,… are the electric field patterns of the fundamental modes on port $i$ 1, 2, 3, … , and we assume that the fields are normalized with respect to the integral of the power flow across each port cross section, then the computed mode field at all output port boundaries is:

$$\boldsymbol{E}_c = \sum_{i=1} S_{i1}\, \boldsymbol{E}_i$$

$$\phi = -\arg(E^{out}) = -\arg(S_{21})$$
$$T = |E^{out}|^2 = abs(S_{21})\text{^}2$$

Where $S_{21} = \sqrt{\frac{Power\ deliverd\ to\ Port\ 2}{Power\ delivered\ to\ Port\ 1}}$

We swept the geometrical parameters of the nano-pillars, till we could find a set of nano-pillar with side lengths $D$, height $h$ and period $p$, which could provide a phase coverage of 2π at an operation wavelength of 478 nm and with an average transmittance higher than 50%. As described in **Supplementary Figure 2** and **Supplementary Video 2**, we selected a set of 28 nano-pillars with square section with a height $h = 520\ \mathrm{nm}$, a period $p = 400\ \mathrm{nm}$ and a side $D$ ranging from $D = 60\ \mathrm{nm}$ up to $D = 340\ \mathrm{nm}$ with a side step of 10 nm.

It is worth to be noticed that we inverted the final sign of output phase extracted from our COMSOL simulation. We are inverting the phase and polarization because in COMSOL the phase convention is based on:

$$\boldsymbol{E}^{CM}(x, y, z, \phi) = \boldsymbol{E_0}(x, y, z)e^{-i\,\boldsymbol{\phi}} \quad (2)$$

Thus, while in our formalism and metasurface design an increase of the phase value indicates retardation (delay accumulation) in phase profile, in COMSOL, a phase increase introduces an electrical field anticipation.

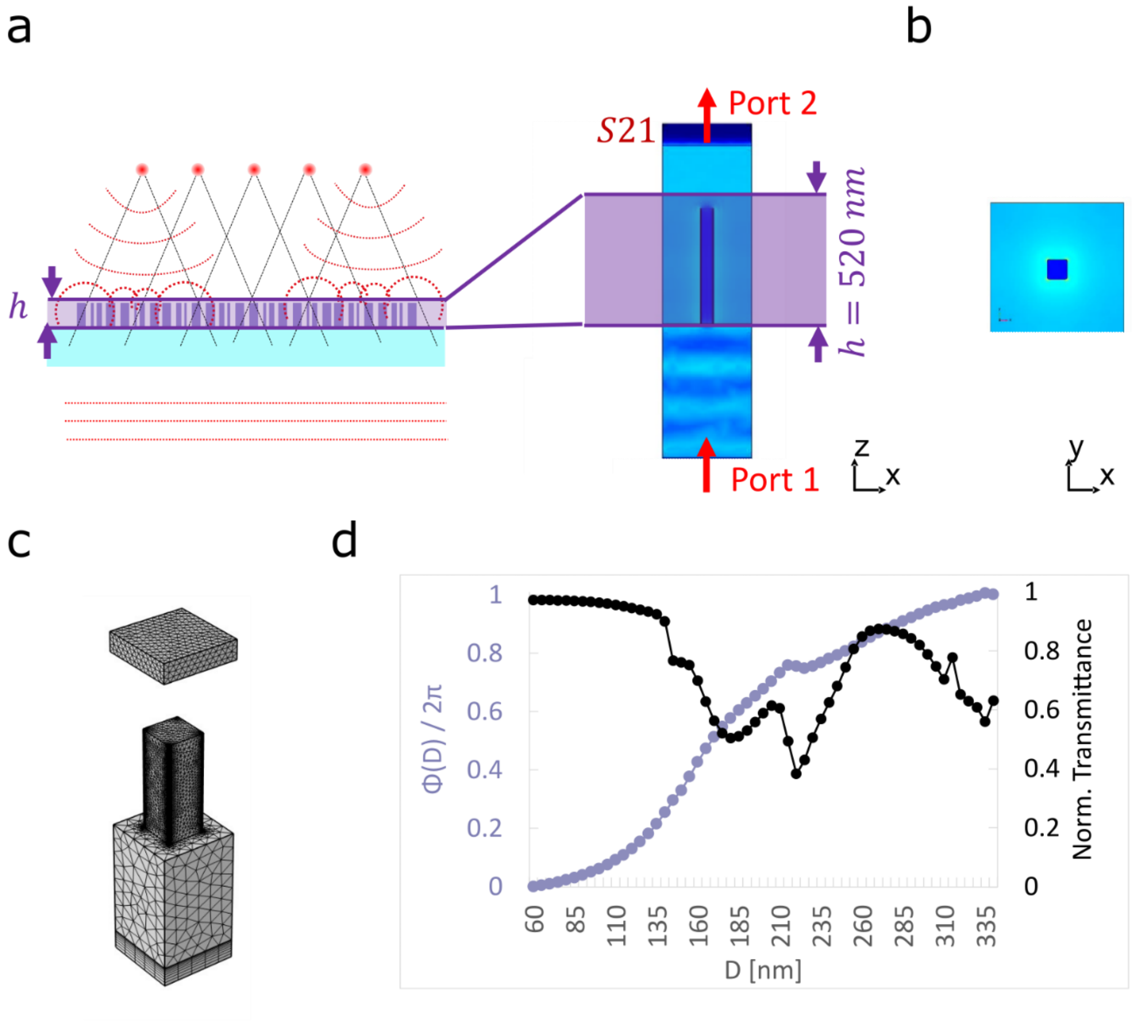


**Supplementary Figure 1 – Numerical simulation of the Silicon Nitride nano-pillar response. a**, Side view of the Silicon Nitride nano-pillar unit (**right** - dark blue) of the metasurface cross section (**left**). Port 1 is the input port at the interface between silicon dioxide substrate and air. Port 2 is the output port on the top of the simulated air volume surrounding the silicon nitride (SiNx) nanopillar structure. The transmittance and phase of the output electrical field are extracted via the scattering parameter $S_{21}$. **b**, Top view of the SiNx nano-pillar unit (dark blue). **c**, 3D view of the mesh generated in COMSOL to simulate the electromagnetic field modulation of each SiNx nano-pillar. **d**, Phase (violet dots) and Transmittance (black dots) of each nano-pillar simulated in COMSOL. The increase of the nano-scatter cross section (diameter D) induces an increase on the relative phase delay accumulated by the incoming light.

***S3. Metasurface design and numerical simulation of the metasurface beam propagation***

The design of our metasurface (MS) is based on a semi-automatized routine written in MATLAB. As described in **Supplementary Figure 3** the code is structured in four main steps. In the first step, the software loads the COMSOL input data and computes the analytical phase profile and numerical aperture (NA) for a metasurface characterized by the input parameters (e.g. focal length, radius, period) set by the user. The two-dimensional (2D) phase profile is then digitalized (second step) where the different nanopillar geometries are distributed over the MS surface. At this point (third step) the software generates the MS layout and computes some quality parameters such as the error. In the last step, we implemented a Beam Propagation Method (BSM) to retrieve the actual MS beam profile and to extract the MS focus position, its FWHM, its depth of focus and its efficiency.

**Supplementary Figure 4** shows the MS layouts of a MS for BB LLS generation obtained with the Silicon Nitride nano-pillar geometries described in **Supplementary Note 2**. The layout is obtained starting from a phase profile of a convergent lens with focal length f = 2000 µm and diameter of 200 µm, for an operating wavelength of 478 nm.

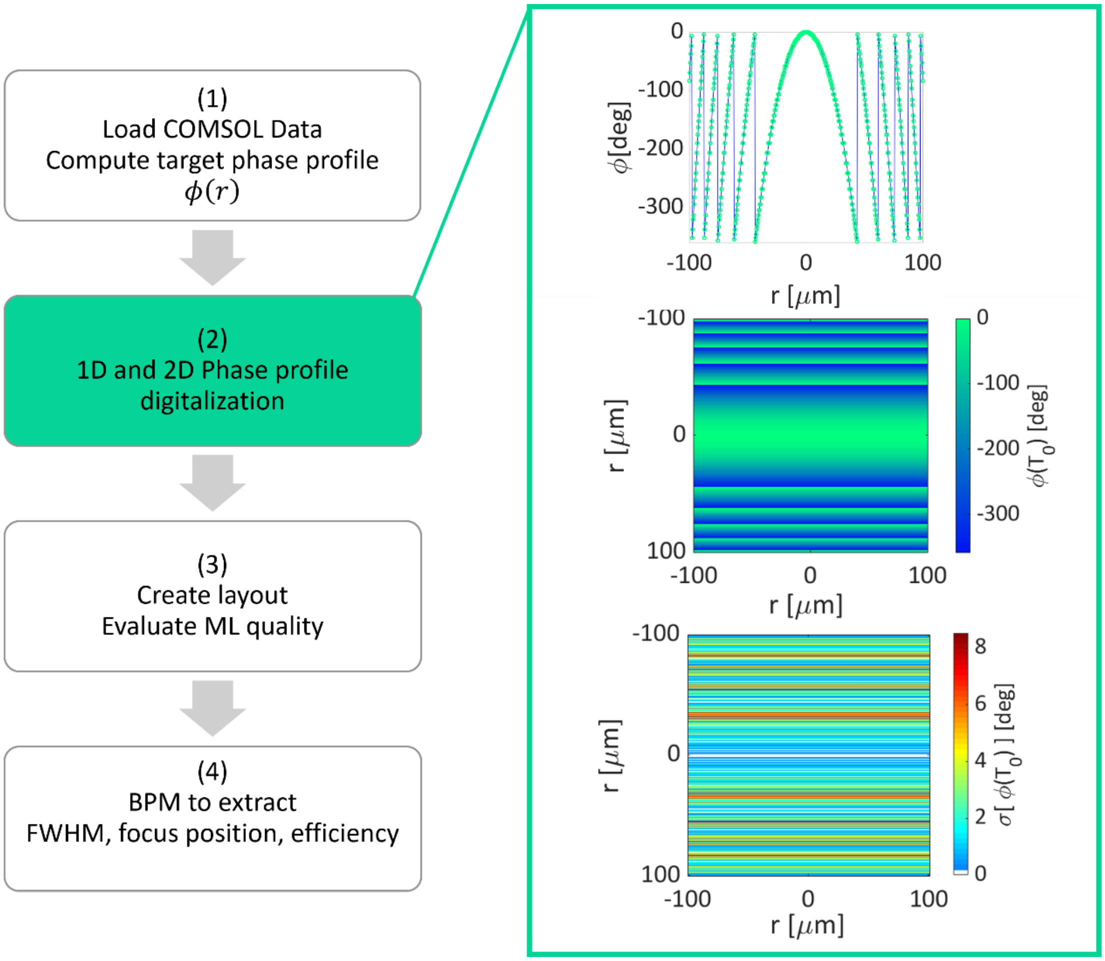


**Supplementary Figure 2 – Metasurface phase design. – 1D and 2D visualization of the MS design quality compared to theoretical values. Left,** Main step of the MS design routine. First (1) the phase and transmittance of each nano-pillar simulated with COMSOL are loaded. Secondly (2), the one-dimensional (1D) and two-dimensional (2D) target phase profiles are generated and digitalized with the discrete parameter space of the nano-pillar phase. Then (3) the metasurface layout is generated and the error on the phase is calculated and displayed. Finally (4) the beam propagation is simulated and displayed with its relative quantitative information. **Right,** an example of the plots generated by the second step of the pipeline: the digitalized 1D phase profile (top) where the circle markers represent the actual MS phase values at a specific radial position overlapped to the theoretical phase shift values displayed with a continuous black line; the 2D phase profile generated with our nano-pillar library (center); the error on the phase, i.e. distance from the theoretical value (bottom).

a

b

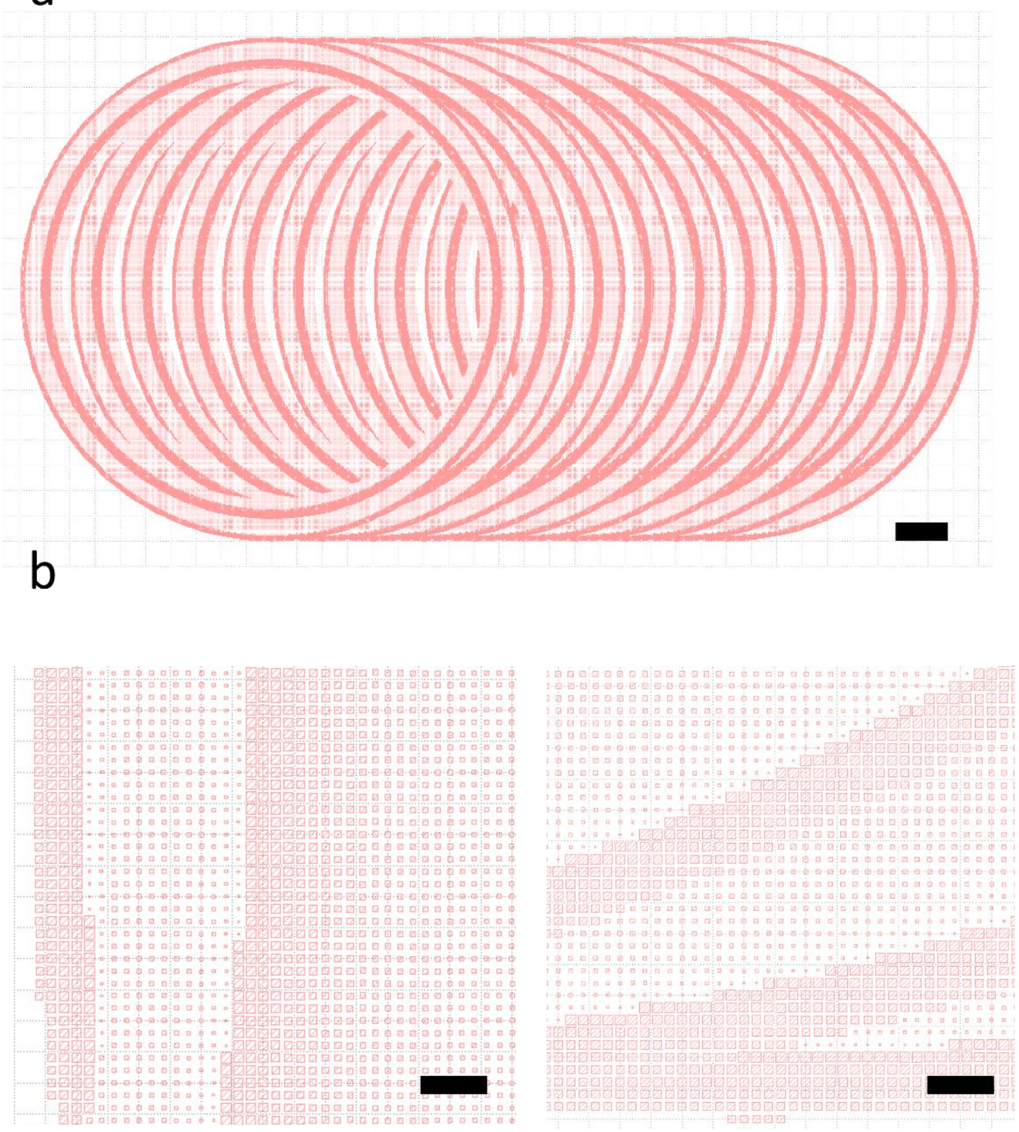

**Supplementary Figure 3 – Sketch of the Bessel beam lattice light sheet (BB-LLS) metasurface layout**. **a**, BB-LLS mask layout. Scale bar 20 µm. **b,** Two details of the BB-LLS mask layout showing the scaling of the cross section of the nano-pillars required to control the relative phase delay of the incoming light.

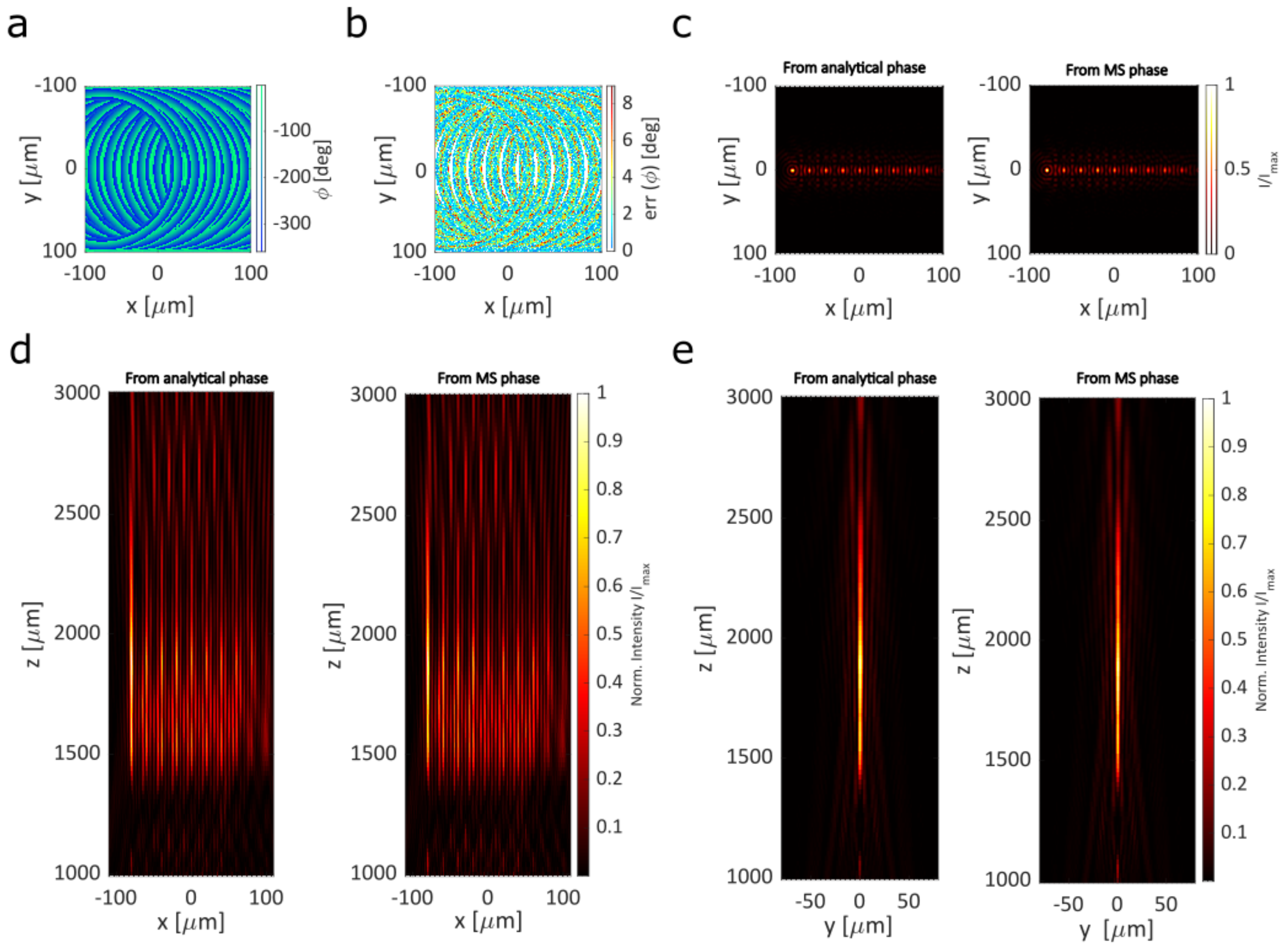


**Supplementary Figure 4 – Beam profile obtained from the ideal phase profile (analytical phase) and from the discretized phase profile of our metasurface achieved with a set of 28 nanopillars.** **a,** Discretized phase profile of our metasurface (MS). **b,** Error on the phase profile. The error is computed as the distance of the discretized phase from the theoretical phase. **c,** Simulation of the XY beam profile of the BB-LLS generated with an ideal phase profile (analytical result – **left**) and with our metasurface phase profile (**right**). **d,** Simulation of the XZ beam profile of the BB-LLS generated with an ideal phase profile (analytical result – **left**) and with our metasurface phase profile (**right**). **e,** Simulation of the YZ beam profile of the BB-LLS generated with an ideal phase profile (analytical result – **left**) and with our metasurface phase profile (**right**).

***S4. Metasurface fabrication and characterization***

We fabricated our metasurface in the clean rooms of the Center of MicroNano Technology CMi at EPFL. The fabrication of our Silicon Nitride metasurfaces requires e-beam lithography technology to achieve up to 5 nm resolution. **Supplementary Figure 5** and **Supplementary Table 1** describe the main steps of the fabrication process. We inspected the metasurface chip surface with both optical and scanning electron microscopy as shown in **Supplementary Figure 6** and **Supplementary Figure 7**.

We systematically performed our MS characterization using a semi-automatized procedure with the setup described in **Supplementary Figure 8**. The stack of images of the beam profile focused by the MS were acquired using µManager[1] to control both the motorized z-stage and the camera. We then reconstructed the beam profile with a MATLAB-based script that we wrote to retrieve the main parameters of the beam.

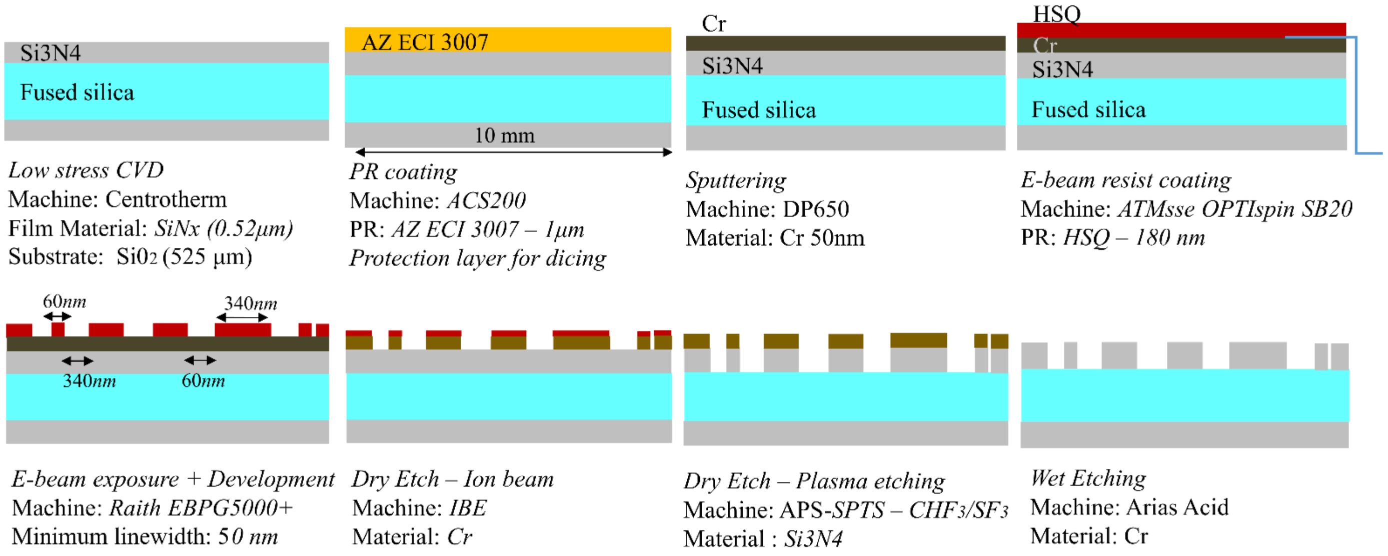


**Supplementary Figure 5 – Metasurface fabrication process flow**. A 520 nm thick Silicon Nitride film is deposited with a Chemical Vapor Deposition process (CVD) on a 525 µm-thick Silicon dioxide substrate. A 1.5 µm thick AZ ECI 3007 resist film is spin coated on the SiNx surface as protective layer for the wafer dicing step. After a cleaning step to remove the resist, a Chromium (Cr) layer is spattered as both reference layer for the EBAM machine and as hard mask. About 200 nm of hydrogen silsesquioxane (HSQ) is coated as positive resist for the e-beam lithography. The desired MS pattern appears on the photoresist after development. The lens structure is then transferred into the Cr and $SiN_x$ layer by one first step of ion beam etching (IBE) followed by a reactive plasma etching step. Finally, the residual of Cr and photoresist is stripped by wet acid etching.

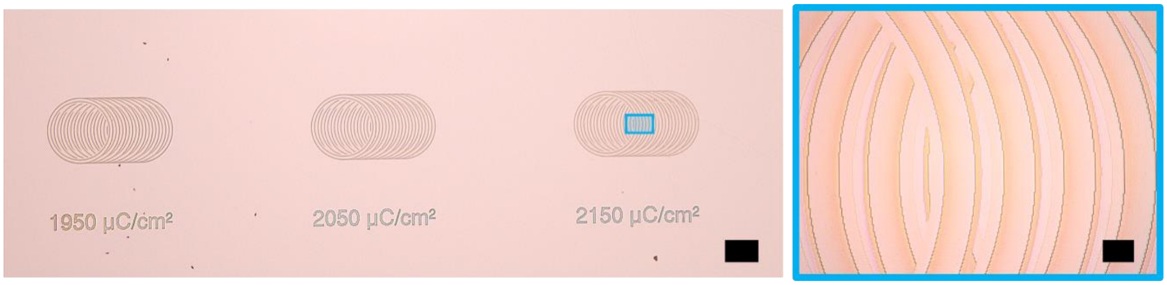


**Supplementary Figure 6 – Inspections of the metasurface layouts after e-beam lithography and HSQ development.** Three metasurfaces (MSs) for Bessel beam LLS generation. Focal length 2000 µm. Scale bar: 100 µm (left); 10 µm (right).

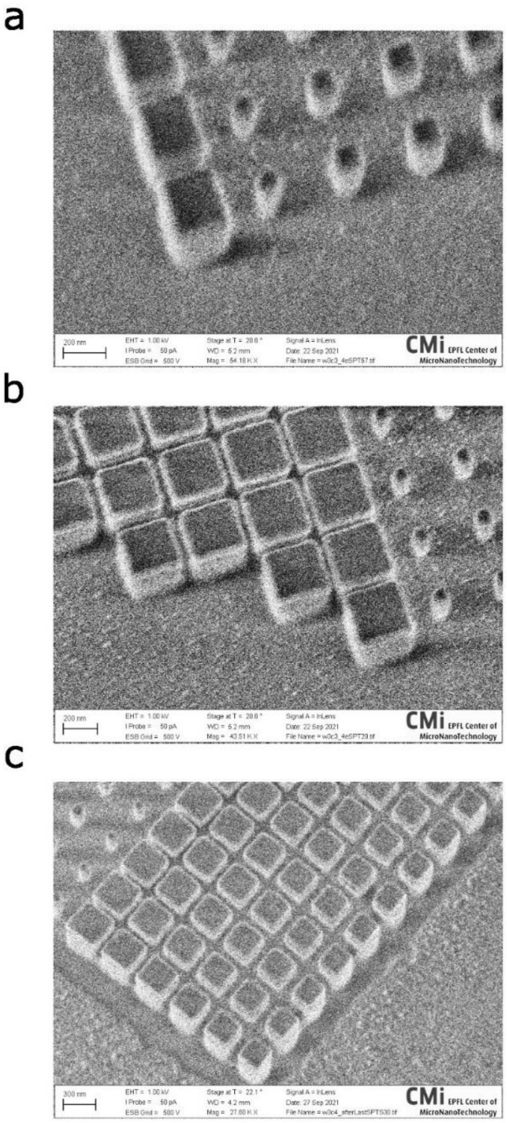


**Supplementary Figure 7 Scanning electron microscopy (SEM) inspections of the metasurface nanopillars. a**, and **b**, Scale Bar 200 nm. **c**, Scale Bar 300 nm.

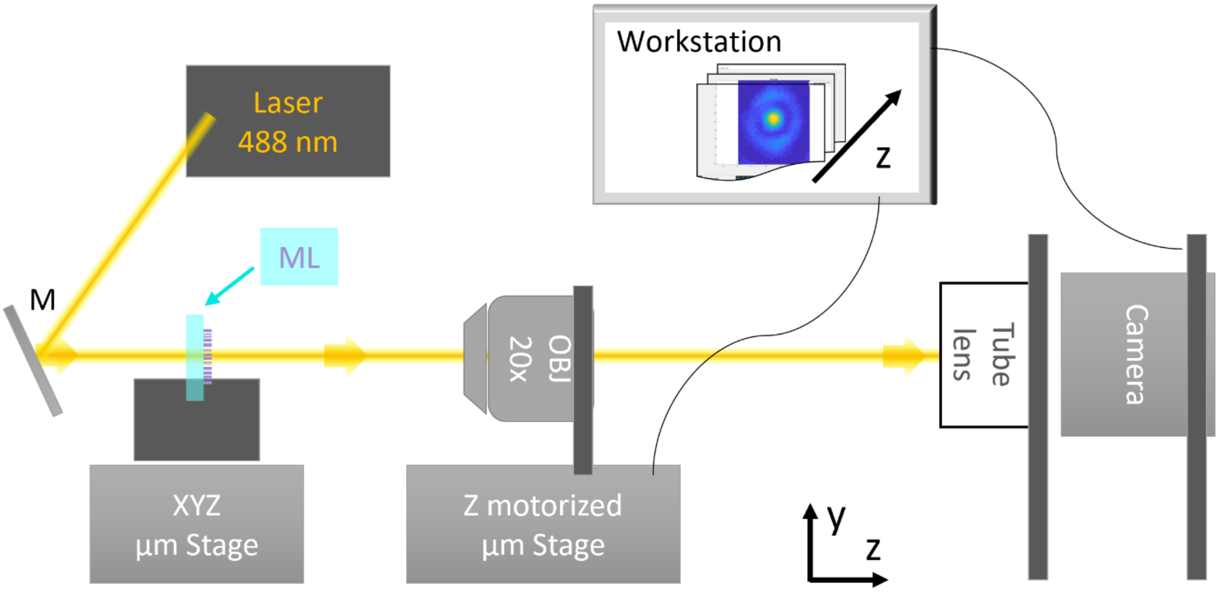


**Supplementary Figure 8 – Sketch of the metasurface characterization setup.** The laser beam is aligned perpendicular to the metasurface (MS) plane. The MS is fixed in the vertical slit of a custom-made holder mounted on a xyz-stage. The detection side is composed by: the 20x Olympus objective (NA = 0.4), the tube-lens (focal length f = 200 mm) and the camera (XIMEA, MQ013RG-ON) connected to the acquisition workstation. The objective is mounted on a motorized stage to automatically acquire the beam profile propagating along the axial (z) direction. Experimental parameters: laser wavelength λ = 488 nm; laser power P = 0.2 mW; camera exposure time Exp = 1 ms; back projected pixel size = 216 nm.

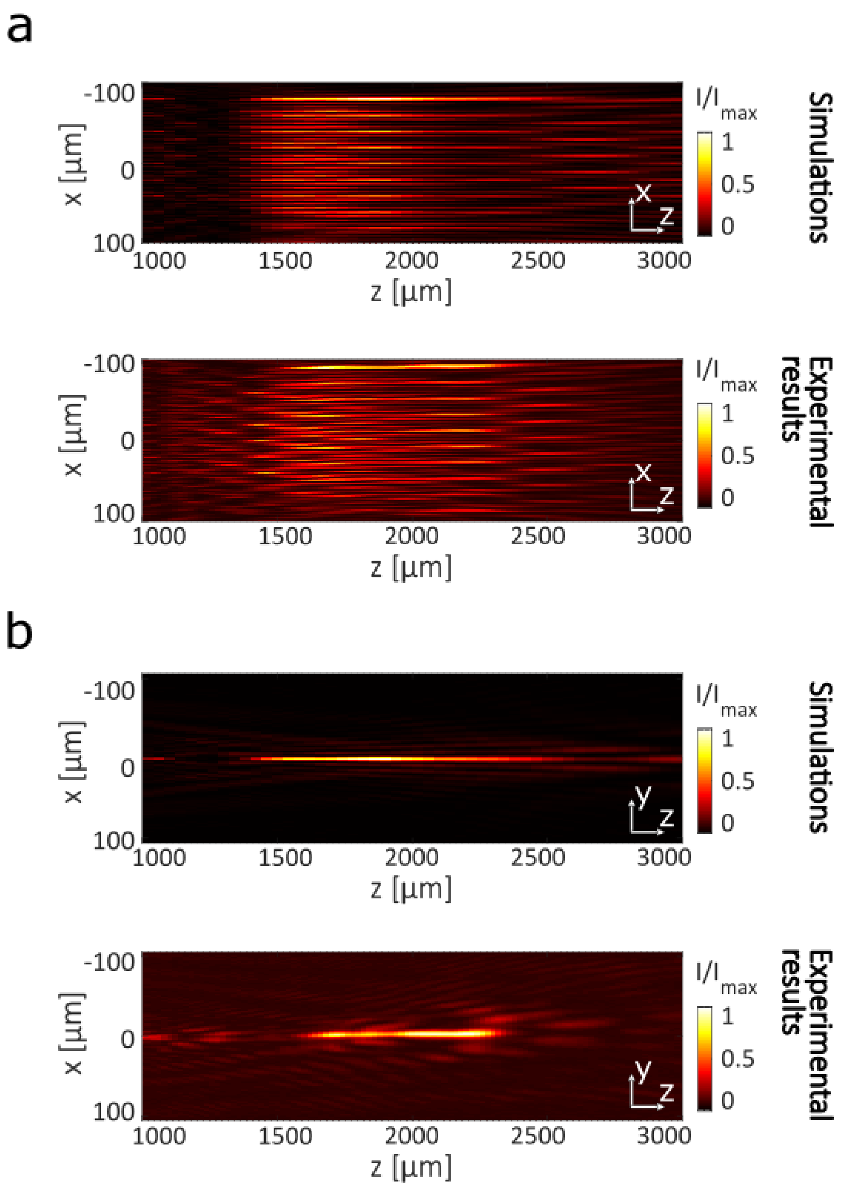


**Supplementary Figure 9 – Metasurface based Bessel beam lattice light sheet: simulation and characterization. a,** Simulation of the YZ beam profile of the BB-LLS generated by our metasurface (MS) design (**top**); experimentally measured beam profiles of the BB-LLS generated by our fabricated MS (**bottom**). **b,** Simulation of the YZ beam profile of the BB-LLS generated by our metasurface (MS) design (**top**); experimentally measured beam profiles of the BB-LLS generated by our fabricated MS (**bottom**). Experimental parameters: laser power P < 0.5 mW; laser wavelength λ = 488nm; camera exposure time Exp = 1 ms.

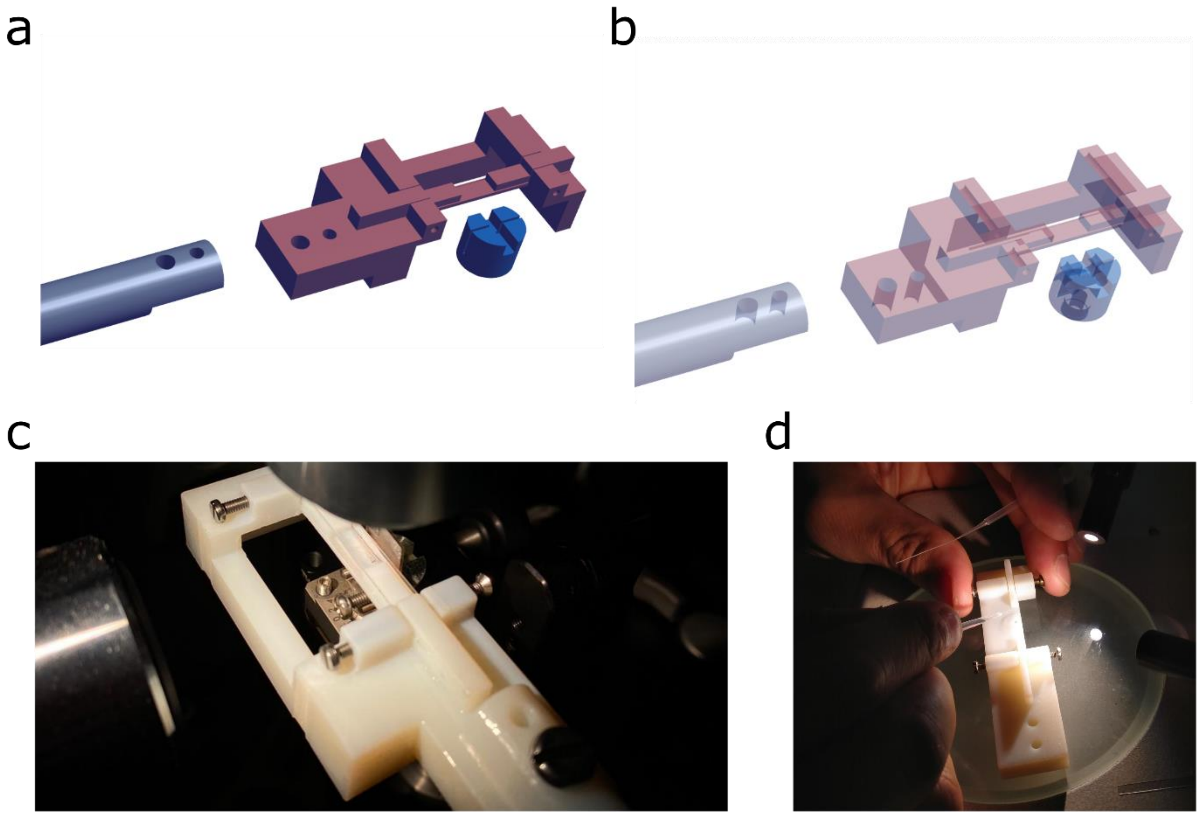


**Supplementary Figure 10 – Custom made holder designs for MS-chip mounting and zebrafish imaging. a,** 3D solid views of the metasurface chip holder (dark-blue) and the 3D-printed holder for zebrafish embedding (brown). **b,** 3D transparent views of **a**. **c,** 3D-printed zebrafish holder with the zebrafish embedded in agarose within the imaging well. The edges of the zebrafish well are 170 um thick glass slices cut with a rectangular shape (3 mm x 40 mm) and inserted in the corresponding slits engraved in the 3D-printed holder. **d,** One of the step of the zebrafish embedding process: the sagittal plane of the zebrafish is manually oriented parallel to the metasurface chip immediately after placing the zebrafish larva within the agarose drop and before the agarose hardens.

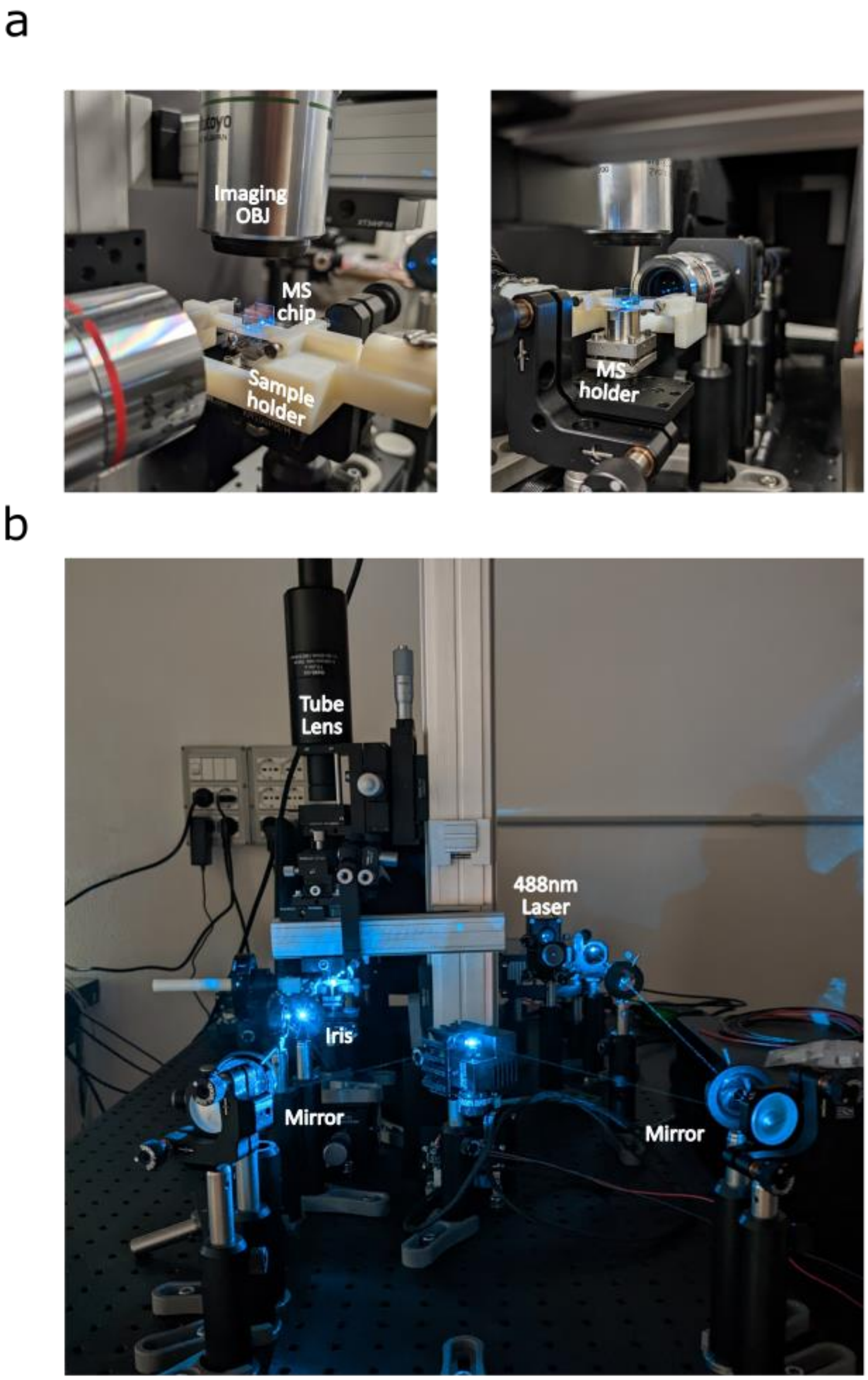


**Supplementary Figure 11 – MS-based Bessel beam lattice light sheet microscope.** Photos of our custom metasurface holder and zebrafish holder (**top**) and of our MS-based BB-LLS setup (**bottom**).

### SUPPLEMENTARY TABLES

#### *Fabrication process*

| Step | Process description |
|---|---|
| 01 | SiNx low-stress LPCVD deposition<br>(520 nm - metasurface layer) |
| 02 | Cr sputtering<br>(40 nm) |
| 03 | HSQ coating<br>(1700 rpm, 180nm) |
| 04 | E-beam lithography<br>(Dose: 1950:2150 uC/cm$^2$, 1 nA, 33MHz, @100 kV) |
| 05 | HSQ development<br>(3 min) |
| 06 | Cr etching (mask)<br>(30 sec) |
| 07 | SiNx plasma etching (metasurface realization)<br>(SPTS APS, $CHF_3/SF_6$ chemistry; etch time: 12 min) |
| 08 | Cr wet etching<br>(Cr_01_UN3264, 3 min) |

**Supplementary Table 1** Chip fabrication process: main steps

#### ***Experimental condition and localization results***

<table>
<tr><th>Sample/ Figure</th><th>Exposure time [ms]</th><th>Laser λ [nm]</th><th>Laser nominal [mw]</th><th>Laser at the sample [mW]</th><th>Number of Frames</th><th>Raw Image pixel size [nm]</th><th>Analysis Algorithm</th></tr>
<tr><td>Fig.2</td><td>1</td><td>488</td><td>22mA/0.5 mW</td><td>-</td><td>200</td><td>216</td><td>Custom MATLAB routine</td></tr>
<tr><td>Fig.3a</td><td rowspan="4">500</td><td rowspan="4">488</td><td rowspan="4">60</td><td rowspan="4">1</td><td rowspan="4">400</td><td rowspan="4">240</td><td rowspan="4">Suite2P[2]</td></tr>
<tr><td>Fig.3b</td></tr>
<tr><td>Fig.3c</td></tr>
<tr><td>Fig.3d</td></tr>
</table>

**Supplementary Table 2** Experimental condition and dataset analysis information.
